# Histone H2A Ubiquitination Mediates the Establishment of Reactivation-Competent HSV-1 Latent Infection

**DOI:** 10.1101/2025.11.23.690028

**Authors:** James Boehlke, Sara A. Dochnal, Chandra Shekhar Misra, Alison Francois, Chris Boutell, Anna R. Cliffe

## Abstract

Herpes simplex virus 1 (HSV-1) establishes a latent infection of neurons, where it persists for life with the capability to reactivate. During latent infection, heterochromatin is deposited onto the viral genome to presumably permit viral gene silencing. Repressive histone modifications associated with the latent genome include the constitutive heterochromatin modifications, H3K9me2/3, and the facultative heterochromatin modification, H3K27me3. The role of these different types during entry into latency and as substrates for reactivation is unknown. H3K27me3 is associated with Polycomb silencing, and a second modification also associated with Polycomb silencing, especially in pluripotent cells and during early development, is H2AK119ub1. Here we found that H2AK119ub1 is enriched on a sub-population of latent HSV genomes. We examined the contribution of both H2AK119ub1 and H3K7me3 deposition to HSV-1 gene silencing during entry into latency. We found that H2AK119ub1 was deposited prior to H3K27me3 and that only inhibition of H2AK119ub1 deposition plays a subtle role in latency establishment. Importantly, we found that inhibiting the enzymatic activity of Polycomb Repressive Complex 1 (PRC1) that deposits H2AK119ub1 during latency establishment prevented later reactivation. In contrast, inhibiting the activity of PRC2, which deposits H3K27me3, did not impact reactivation. Together, these data demonstrate the heterogenic nature of the epigenetic structure of latent HSV genomes and provide evidence that those associated with H2AK119ub1 are the template for reactivation.

## Introduction

Human herpesviruses establish lifelong latent infections in specific host cells. During latency, viral genomes are typically associated with repressive heterochromatin, which likely maintains long-term gene silencing and enables viral persistence. However, heterochromatin is highly diverse and can vary in the mechanisms of formation, the fate of the chromatin structure, and its reversibility. Because herpesvirus genomes periodically reactivate from latency to produce infectious progeny, they must remain transcriptionally silent yet retain the capacity to revert to euchromatin and support lytic gene expression. As highly evolved viruses, herpesviruses have likely coevolved with their host species to exploit certain types of heterochromatin that permit such transitions. Therefore, investigating heterochromatin-based silencing mechanisms on herpesvirus genomes provides insights into the host mechanisms of gene silencing and reversal, and also informs strategies to prevent viral reactivation.

Herpes Simplex Virus-1 (HSV-1) establishes a latent infection in post-mitotic peripheral neurons, and reactivation can cause lesions, recurrent pain, blindness, and encephalitis. There is also accumulating evidence linking HSV-1 infection with late-onset Alzheimer’s disease (1). The latent HSV-1 viral genome is known to be associated with heterochromatin, with the enrichment of certain histone post-translational modifications (PTMs) including H3K27me3, which is indicative of Polycomb silencing (2–5), as well as H3K9me3 (3), which is characteristic of more long-term constitutive heterochromatin. Although associated with repressive histone modifications, it is currently unknown whether heterochromatin deposition on the viral genome directly drives gene silencing. An alternative is that heterochromatin is deposited on already silenced genes to maintain long-term gene silencing and viral latency. This is important to understand to gain insight into how the virus establishes a latent infection of neurons.

EZH1 and EZH2 catalyze the addition of K27 methylation onto histone H3 as part of the Polycomb repressive complex 2 (PRC2) (6). Previously, PRC2 has been found stably associated with the HSV-1 genome as latency is established, but only after lytic gene silencing had already occurred (2), indicating that PRC2 activity may not directly drive lytic gene repression during entry into latency. On the cellular genome, H3K27me3 can be accompanied by the mono-ubiquitination of H2AK119 (H2AK119ub) (7). H2AK119ub is deposited by the Polycomb Repressive Complex 1 (PRC1) (8–10). Although H2AK119ub was initially thought to be deposited after H3K27me3, multiple recent studies have found that H2AK119ub is deposited first, followed by H3K27me3, at least in embryonic stem cells and during early development (11–14). In this context, H2AK119ub deposition by PRC1 but not H3K27me3 deposition drives gene silencing. H3K27me3 can form in the absence of H2AK119ub1, and this may play a more important role in maintaining gene silencing, often referred to as the primary form of Polycomb silencing in terminally differentiated cells (15–17). HSV-1 establishes a latent infection in non-mitotic, terminally differentiated neurons, where the contribution of H2AK119ub1 to driving gene silencing is unknown. Therefore, the association of H2AK119ub with HSV-1 is also unknown. Further, when the HSV-1 genome enters a host cell nucleus, it does so as epigenetically naïve, meaning it does not contain a pre-existing chromatin template (18, 19). Therefore, HSV-1 also serves as a tool to study the basic mechanisms of Polycomb recruitment and silencing in terminally differentiated neurons.

Understanding the contribution of heterochromatin silencing to HSV latency and reactivation requires considering that epigenetic structures across individual genomes may be heterogeneous, with different types of heterochromatin playing distinct roles in the initiation of gene silencing. HSV-1 latent infection is heterogeneous, with altered expression of the latency-associated transcript (LAT) and differential localization of viral genomes to distinct subnuclear regions (20). Further, at any one time only a fraction of viral genomes undergo reactivation (21–25). We recently reported that the association of HSV-1 genomes with H3K9me3 and the reader protein ATRX results in a form of latent infection that is restricted for reactivation (26). However, it is currently unknown whether there is a specific HSV-1 epigenetic structure that is more reactivation-competent.

Because of its reversible nature, Polycomb silencing is a key candidate for reactivation competent HSV-1 genomes (27). Accumulating evidence suggests that H2AK119ub marks regions on the host genome that are silenced during early stages of development, with the capacity to transition to euchromatin for gene expression upon differentiation (28). Therefore, here we set out to determine whether H3AK119ub was even associated with the latent viral genome. We found enrichment of this modification on latent viral genomes. Further, we found that inhibiting the deposition of H2AK119ub1 during latency establishment had a mild impact on viral gene expression at early times, whereas inhibiting PRC2 activity had a negligible impact. Importantly, though, PRC1 inhibition during latency establishment dramatically reduces the ability of latent viral genomes to reactivate. These data, taken together, support a model in which PRC1 mediates lytic viral gene silencing in a manner amenable to later reactivation.

## Results

### H2AK119ub is enriched on the HSV-1 genome during latent infection established *in vivo*

The latent HSV-1 genome is known to be associated with the H3K27me3. However, whether H2AK119ub can be targeted to a viral genome in terminally differentiated neurons was unknown. Therefore, we initially used a well-established *in vivo* model of HSV-1 latency to determine whether H2AK119ub was detectible on the viral genome (Fig. 1A). We infected mice via the ocular route and at 30 days post-infection, we harvested latently infected trigeminal ganglia (TG) and performed chromatin immunoprecipitation (ChIP) assays for H2AK119ub, as well as negative control IgG as previously described (2). We observed a significant enrichment of H2AK119ub on repressed lytic viral gene promoters during latent infection over IgG controls (Fig 1B), as determined by quantitative PCR (qPCR) with primers specific for the *ICP27* (an immediate-early gene) and *ICP8* (an early gene) promoter regions.

**Figure 1.**
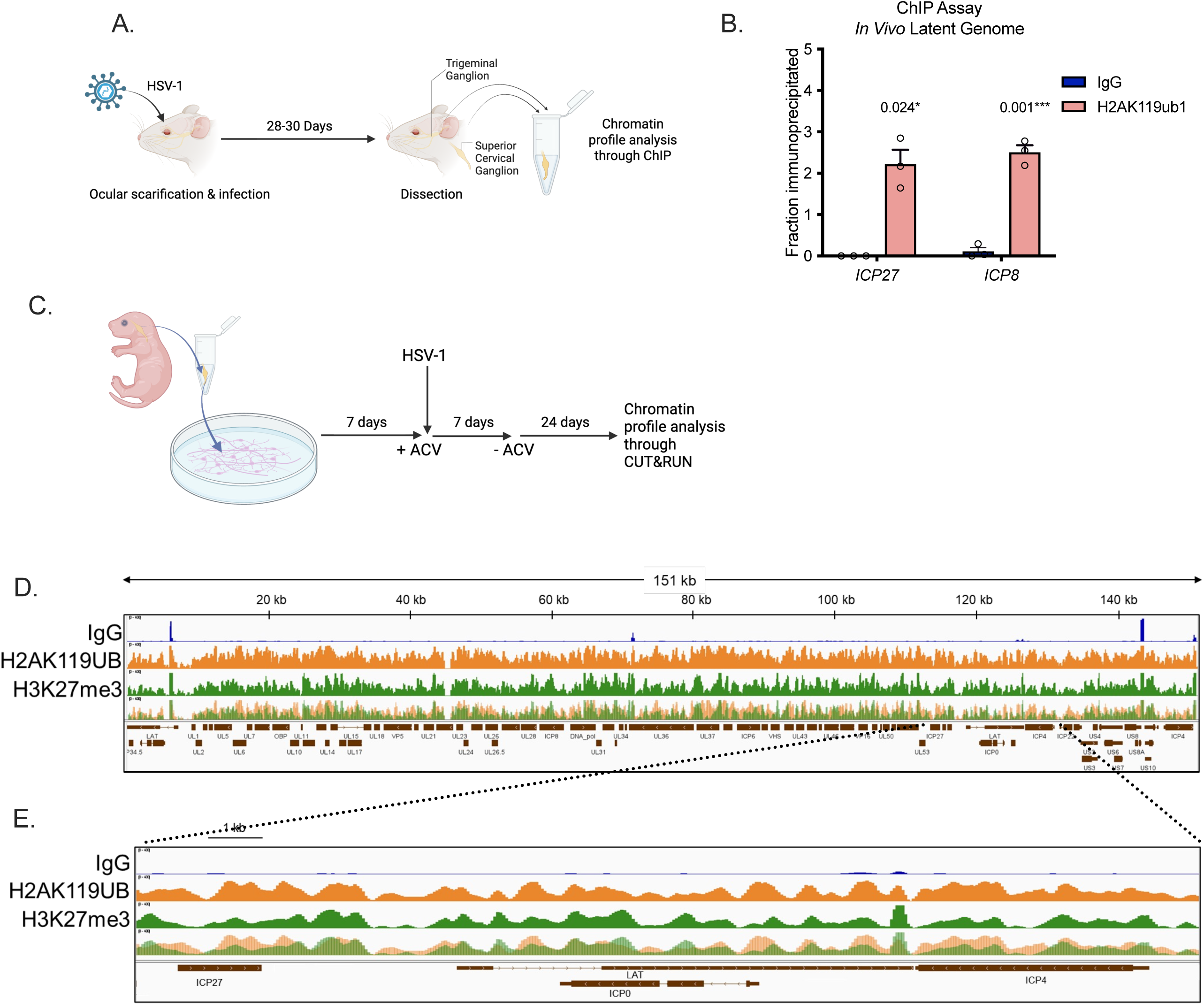
H2Aub1 associates with the latent HSV-1 genome. A) Schematic of in vivo experiments with HSV-1. B) Mice were infected with WT HSV-1, and latent trigeminal ganglia (TG) were collected 30 days post-infection. Chromatin immunoprecipitation (ChIP) was performed using an IgG control or H2AK119ub, and quantitative PCR (qPCR) was performed to analyze binding to select promoters of the immediate-early viral gene ICP27 and the early viral gene ICP8. Enrichment over Input determined. 3 biological replicates; Welch’s t-test. Normality determined by Shapiro-Wilk test. Means and SEMs are represented. *, P□<□0.05; **, P□<□0.01. ns, not significant. C) Schematic of infection of primary neurons with HSV-1 D) Primary neurons were infected with HSV-1 Stayput-GFP in the presence of viral DNA replication inhibitor acyclovir/ACV (10 μM during inoculation period and 50 μM following removal of inoculum). CUT&RUN was performed on neurons collected and permeabilized 30 days post-infection using IgG control (blue), H2AK119ub (orange), or H3K27me3 (green). Raw reads from merged experimental replicates aligned across the HSV-1 genome. E) Zoomed in region as marked. A and C were Created in BioRender. Cliffe, A. (2025) https://BioRender.com/jdycuyb and Cliffe, A. (2025) https://BioRender.com/ta6o69j

### H2AK119ub and H3K27me3 associate with a population of latent HSV-1 genomes *in vitro*

To mechanistically investigate the contribution of Polycomb Group proteins to HSV-1 latency, we turned to an in vitro system where we can more readily manipulate their function solely in neurons. First, we mapped the association of H2K119ub and H3K27me3 on the latent viral genome following neuronal infection in vitro. We infected primary neonatal sympathetic neurons isolated from the superior cervical ganglia of newborn mice with HSV-1 Stayput-GFP (21) at an MOI of 10 PFU/cell in the presence of viral DNA replication inhibitor acyclovir (ACV) to promote latency establishment (Fig. 1C). Neurons were collected 30 days post-infection, a time point at which latency has been fully established *in vitro* (21) and mirrors what is documented *in vivo* (2–4). Primary neurons also have the advantage that they are more amenable to performing Cleavage Under Target & Release Under Nuclease (CUT&RUN), which gives a higher resolution than ChIP and it requires less input material. We performed CUT&RUN-sequencing using control IgG, along with antibodies against H2AK119ub and H3K27me3. Both H2AK119ub and H3K27me3 were broadly enriched over IgG controls, as well as input, across the latent HSV-1 genome, although in more discernable peaks versus the ChIP-Seq data (Fig 1D). Both H2AK119ub and H3K27me3 were enriched on lytic genes and intergenic regions, including but not limited to immediate early genes ICP27 and ICP0 (Fig 1E). Both modifications were but were notably depleted at between CTCF sites in the transcriptionally active LAT promoter region (Fig 1E). Therefore, both H3K27me3 and H2AK119ub are enriched on the latent HSV-1 genomes following latency established either in vivo or in vitro.

### A sub-population of HSV-1 genomes associates with H2AK119ub1 and/or H3K27me3 during latent infection *in vitro*

Collectively, the data form Chip and CUT&RUN demonstrate genome-wide enrichment of H2AK119ub on the repressed HSV-1 genome during latent infection. However, there is known heterogeneity in HSV-1 latency based on prior lytic gene expression, expression of the LAT, and the sub-nuclear localization of viral genomes (20, 22, 26, 29). Therefore, we were interested in determining whether there is heterogeneity in facultative heterochromatin associated with the latent HSV-1 genome. Interestingly, only a subpopulation of neurons containing latent viral genomes will reactivate at any one time (21, 22, 25, 26, 30–32). Therefore, we next used an additional *in vitro* approach to investigate heterochromatin association with the latent viral genome at the single-cell and individual viral-genome levels. We infected primary neurons with an EdC-labelled HSV-1 virus at an MOI of 10 and fixed at 30 days post-infection. Using a combination of click chemistry and immunofluorescence, we analyzed the 3-dimensional co-localization of the latent HSV-1 genome with PRC1 and PRC2-mediated histone modifications H3K27me2, H3K27me3 and H2Aub1. H3K27me2 was including in the analysis as it is also correlated with gene repression and has recently been shown to associate with the HSV-1 genome during early lytic infection of non-neuronal cells (33). In agreement with the CUT&RUN and ChIP data, we observed viral genome foci that colocalized with clusters of H3K27me3 and H2AK119ub (Fig 2A).

**Figure 2.**
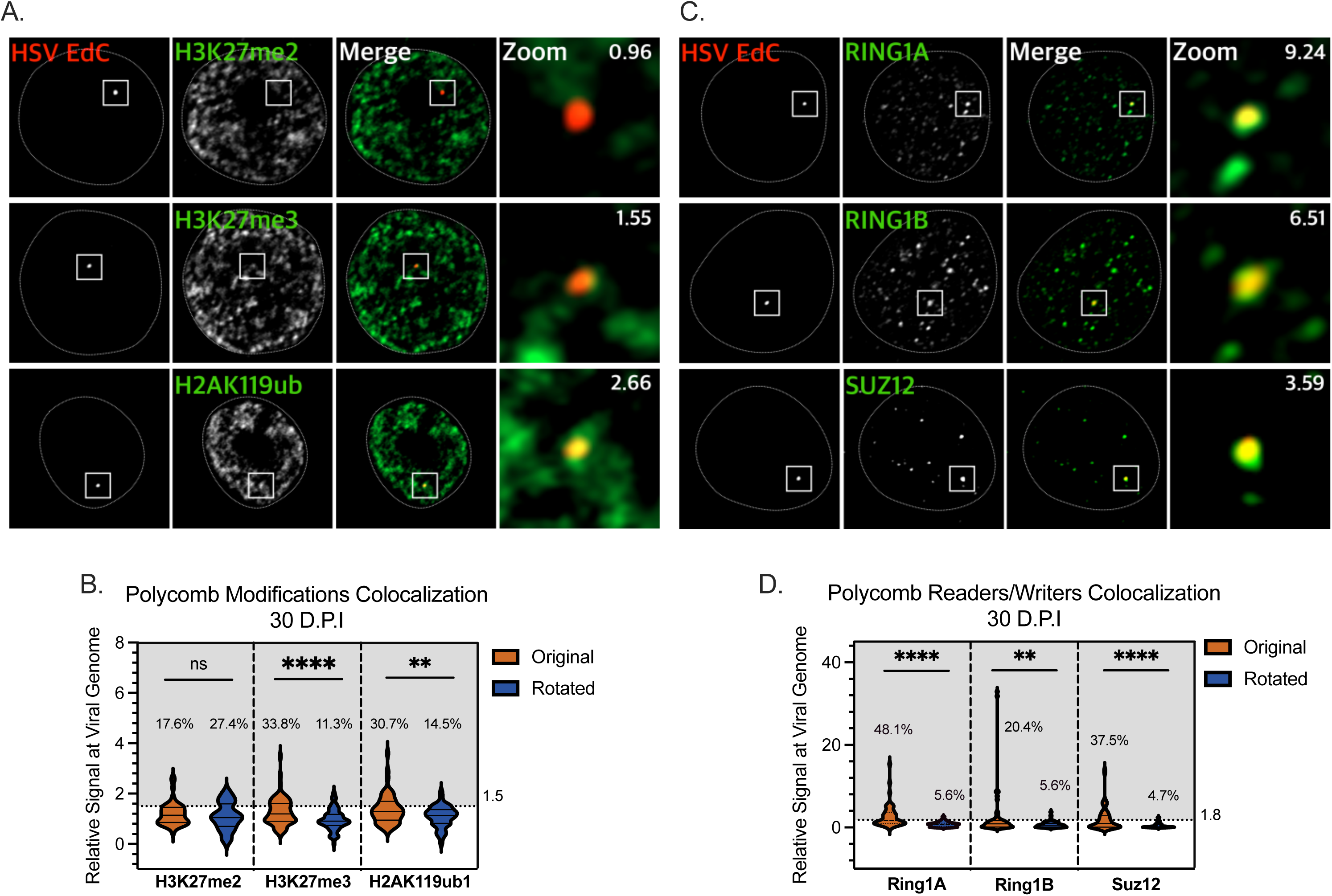
H2Aub1, H3K27me3, and Polycomb Proteins co-localize with a sub-population of HSV-1 genomes during latency. Primary neurons were infected with EdC-labeled HSV-1 in the presence of ACV. 30 days post-infection, latently infected neurons were fixed and stained for Polycomb-associated histone modifications H3K27me3, H3K27me2, or H2AK119ub (A & B) and Polycomb repressor complex 1 (PRC1) components Ring1A and Ring1B, and PRC2 core component Suz12 (C & D). The HSV-1 genome was visualized using Click Chemistry. Enrichment values of chromatin mark/reader/writer with each individual HSV-1 genome analyzed using NucSpotA are shown. B and D) Signal intensity of histone modifications (B) and Polycomb proteins (D) at the viral genome at 30-days post-infection. Each data point represents one viral genome. Percentages indicate the proportion of genomes with NucSpotA intensity ratios above the denoted co-localization threshold (dashed line). Paired comparisons were performed for each image by rotation of the genome channel 90 degrees to achieve random placement. Threshold N>75 cells from 3 biological replicates. Wilcoxon test. Normality determined by Kolmogorov-Smirnov test. *, P□<□0.05; **, P□<□0.01. ns, not significant.

Histone modification stains are broadly nuclear, making it unclear whether overlaps between the green and red signals are a random coincidence or genuine co-localization. Therefore, to measure co-localization in a high-throughput and unbiased fashion, we used our recently developed program NucSpotA (33). NucSpotA measures the enrichment of the modification (based on the intensity of the immunofluorescence signal) at the viral genome relative to the entire nucleus. By also measuring the enrichment values of the same stain on a set of control images where the viral genome is rotated 90 degrees, we can determine whether co-localization with the HSV-1 genome in original images is higher than that obtained for random placement of the viral genome. We further assign a threshold for the enrichment values by eye and calculate the proportion of viral genomes that meet this threshold to estimate what proportion of latent viral genomes are co-localized with these Polycomb modifications and components during latent infection. In agreement with our population-level findings, the HSV-1 genome significantly co-localized with H3K27me3 at 30 days post-infection (Fig 2B). Specifically, we found that H3K27me3 co-localized with a subset of latent viral genomes, approximately 30%. In contrast, only approximately 8% of genomes co-localized with H3K27me2, and this was not significantly enriched at this time point compared to the rotated control (Fig. 2B). Importantly, the latent HSV-1 genome also significantly co-localizes with H2AK119ub in approximately 34% of the infected neurons (Fig. 2B).

We also quantified the co-localization of different PRC1 components with the viral genome, including the core components Ring1A and Ring1B, and the PRC2 component Suz12. We observed staining puncta for all three proteins (Fig. 2C) and viral genes that co-localized with these puncta in approximately 48% of neurons for Ring1A, 20% for Ring1B, and 37% for Suz12. Therefore, using multiple methods, our data demonstrate enrichment for H2AK119ub and H3K27me3 but not H3K27me2 on at least a sub-population of viral genomes, and co-localization with components of the complexes that can both read and write these modifications. For at least one of these (Ring1A), the frequency was higher than either modification alone (48% compared to approximately 30% for H3K27me3 and H2Aub1).

### PRC1 associates with the viral genome during active stages of viral gene silencing

As previously determined by performing ChIP on ganglia in a mouse model, PRC2 (specifically Suz12) and associated H3K27me3 are not enriched on the viral genome until lytic viral genes have already been silenced (2). These observed kinetics suggest that PRC2 may not play an active role in initiated lytic gene silencing in neurons. Because of the known function of vPRC1-medited H2AK119ub in driving gene silencing before H3K27me3 deposition (12, 34–37), at least in embryonic stem cells, we posited that H2AK119ub may be deposited prior to H3K27me3. In addition, the dimethyl form of H3K27me2 can precede H3K27me3 and can silence cellular genes independently of H3K27me3 (38). Therefore, we investigated the kinetics of H3K27me2, H3K27me3, and H2AK119ub following the infection of primary neurons at a single-genome resolution. Several time points were preliminarily surveyed, but here we present the acute time point of 3 days post-infection; a time point at which we have previously observed a steep reduction in lytic viral gene expression during latency establishment (21). In agreement with previous *in vivo* findings, H3K27me3 was not found to be significantly co-localized with latent viral genomes at this 3-day post-infection time point (Fig. 3 A&B). In addition, we did not observe co-localization of H3K27me2 with the viral genome (Fig. 3 A&B). In contrast, we found that 20% of viral genomes were positively co-localized with H2AK119ub and the enrichment was significantly enriched compared to the rotation control images (Fig. 3A&B). The core PRC1 components Ring1B and Ring1A, were also significantly co-localized at the early three-day post-infection timepoint (Fig 3. C& D). These data indicate that vPRC1- mediated H2Aub1 deposition on the viral genome occurs prior to PRC2-mediated H3K27me3.

**Figure 3.**
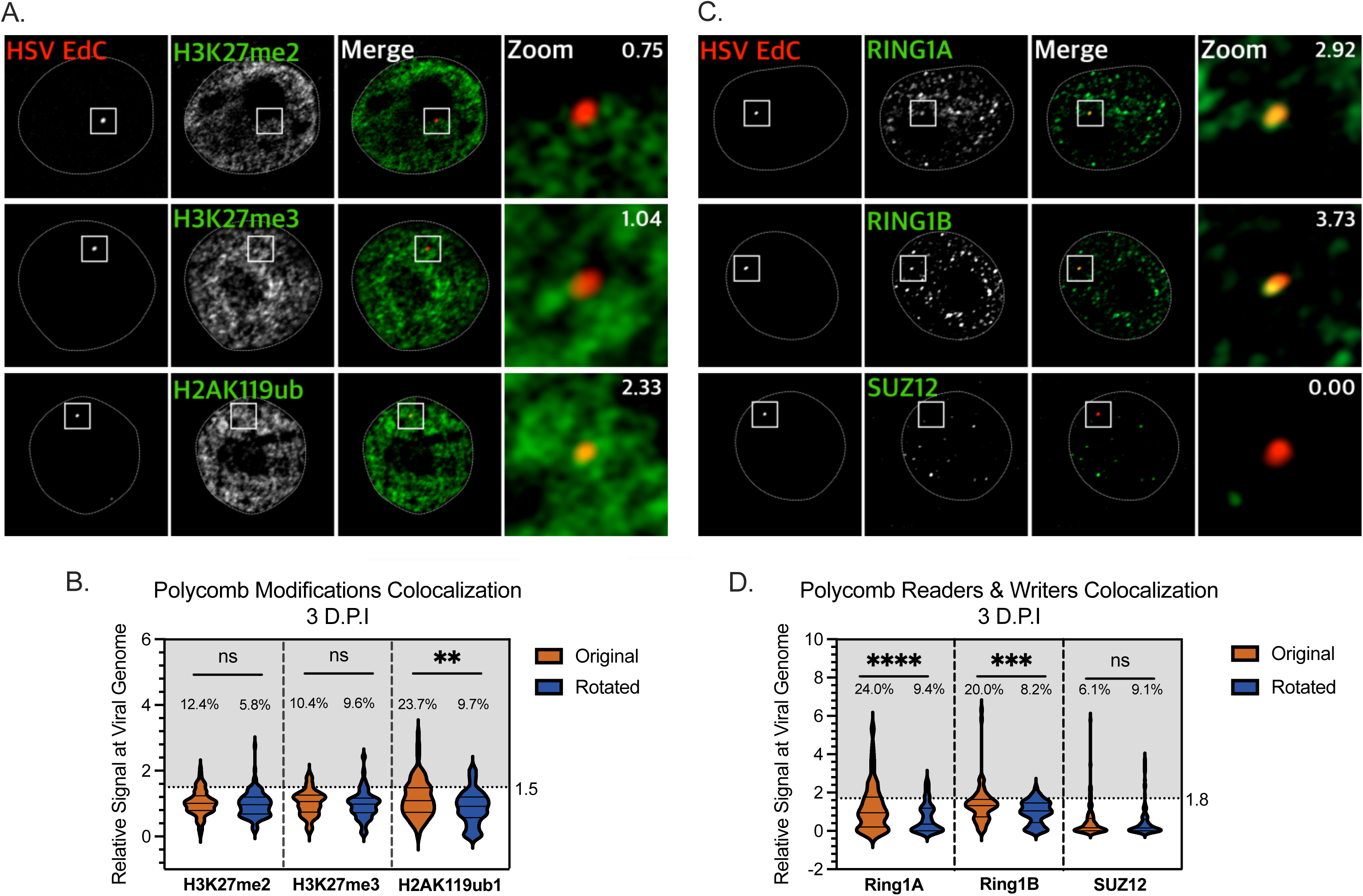
Single-viral-genome analysis demonstrates that HSV-1 genomes associate with PRC1-mediated H2AK119ub during active viral gene silencing. A and B) Primary neurons were infected with EdC-labeled HSV-1 in the presence of ACV. 3 days post-infection, neurons were fixed and stained for histone modifications H3K27me3, H3K27me2 and H2AK119ub (A & B), and PRC1 core components Ring1A, Ring1B and PRC2 core component Suz12 (C & D). The HSV-1 genome was visualized using Click Chemistry. Enrichment values of chromatin mark/reader/writer with each individual HSV-1 genome are shown. 10 μm scale, 1 μm scale for zoom. Signal intensity of histone modifications (B) and Polycomb proteins (D) at the viral genome at 30-days post-infection. Each data point represents one viral genome. Percentages indicate the proportion of genomes with NucSpotA intensity ratios above the denoted co-localization threshold (dashed line). Paired comparisons were performed for each image by rotation of the genome channel 90 degrees to achieve random placement. Threshold N>75 cells from 3 biological replicates. Wilcoxon test. Normality determined by Kolmogorov-Smirnov test. *, P□<□0.05; **, P□<□0.01. ns, not significant.

### PRC1 but not PRC2 inhibition or depletion results in a modest increased in lytic gene expression during acute infection of neurons

PRC1 association with repressed HSV-1 genomes, and the kinetics of this association, suggests PRC1 may actively silence these genomes. However, our data on binding, co-localization, and viral gene repression are thus far only correlative in nature. We therefore took a functional approach to investigate the contribution of PRC1 and PRC2 to the repression of viral lytic gene expression. We infected primary neurons with HSV-1 Stayput-GFP in the presence of PRC1 and PRC2 inhibitors. We used PTC-209 (39) and RB-3 (40), which are both Ring1 inhibitors. PTC-209 has been demonstrated to reduce H2AK119ub on cellular DNA in neurons, although the exact mechanism of PTC-209 is unclear. RB-3 blocks the association of both Ring1A and Ring1B with chromatin and, therefore, the ability to deposit H2AK119ub (40). We also used the PRC2 inhibitors UNC1999, CPI-1205 and EPZ438. We observed a subtle but statistically significant increase in all classes of viral transcripts upon PRC1 inhibition with both PTC-209 and RB-3 (Fig. 4A, Supplemental Figure 1A and 1B). In contrast, the PRC2 inhibitor-treated samples did not display a statistically significant enhancement in the expression of immediate early, early, or late genes surveyed (Fig. 4A and Supplemental Figure 1A and 1B). We verified the loss of H2AK119ub or H3K27me3 by quantifying the genome co-localization in the presence of the PRC1 inhibitor RB-3 or the PRC2 inhibitor EPZ6438. We found that RB-3 significantly reduced the proportion of genomes associated with H2Aub1 from 26 to 17% positive co-localization, which is a number similar to that observed for genome co-localization by chance (Fig. 4B and Supplemental Figure 1C).

**Figure 4.**
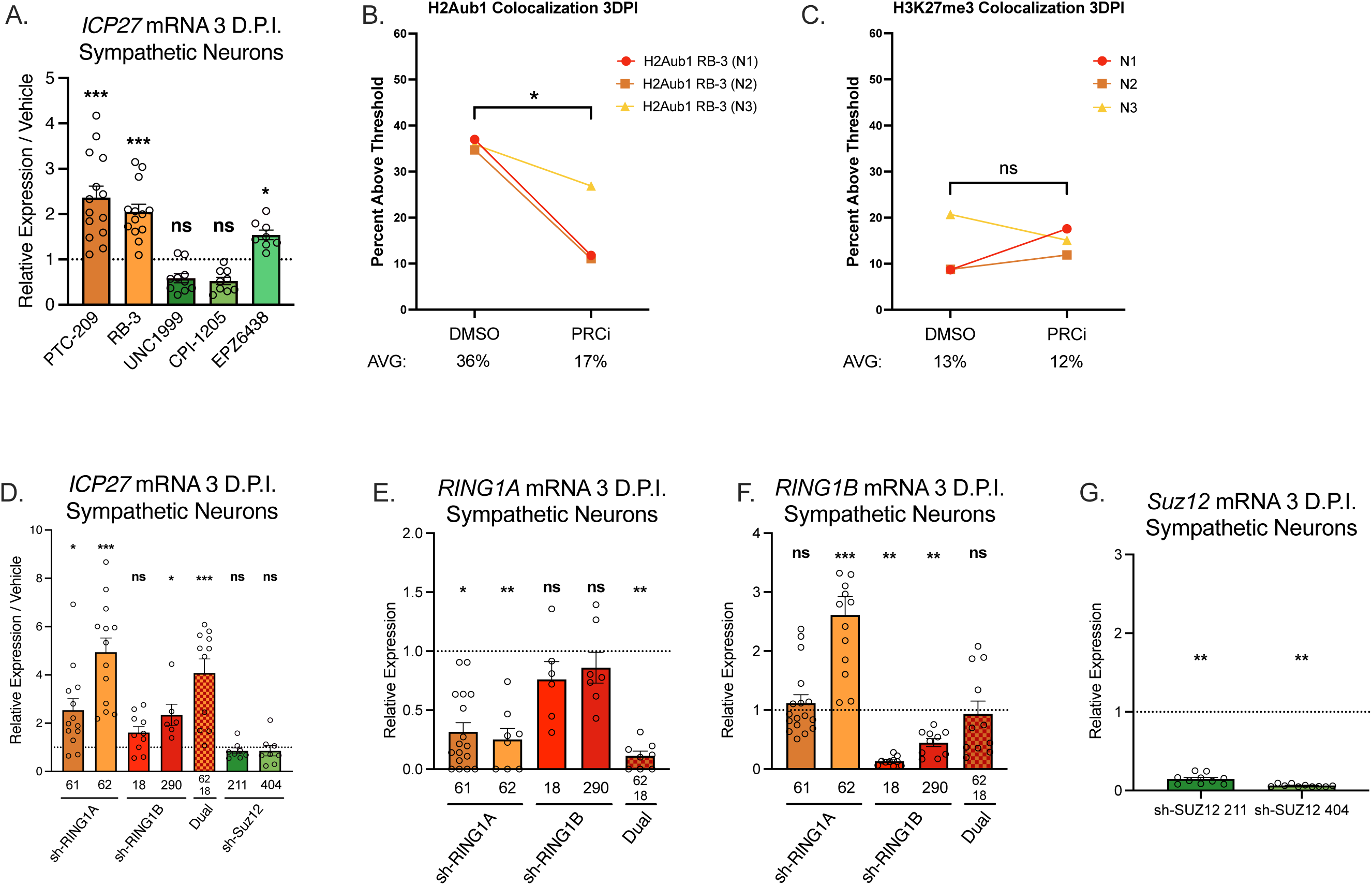
PRC1 but not PRC2 mildly promotes HSV-1 lytic gene silencing during the establishment of latent infection. (A-C) Primary neurons were treated with PRC1 inhibitors PTC-209, RB-3, UNC1999, CPI-12-5 or EPZ6438 pre-infection for 1 hour, during inoculation with HSV-1 Stayput-GFP in the absence of ACV and following the removal of inoculum for 3 days post-infection. Expression of lytic viral immediate early gene ICP27 (A), 3 biological replicates, Wilcoxon Test. B) Quantification of genome co-localization with H2Aub1 following RB-3 treatment. C) Quantification of genome co-localization with H3K27me3 following EPZ6438 treatment. D-G) Neonatal sympathetic neurons were transduced with a non-targeting shRNA control lentivirus or one of two independent lentiviruses expressing shRNAs that target Ring1A (sh-Ring1A 61, sh-Ring1 62) or Ring1B (sh-Ring1 18, sh-Ring1 290), or both Ring1A and Ring1B (sh-Ring1 62 and sh-Ring1 180), or Suz12 (sh-Suz12 211, sh-Suz1 404). Following depletion, primary neurons were infected with Stayput-GFP at a multiplicity of infection (MOI) of 5 in the absence of ACV. (D-G) Relative viral expression at 3 days post-infection compared to shRNA control quantified by RT-qPCR for viral genes ICP27 (D), and cellular Ring1A (E), Ring1B (F) or Suz12 (G) normalized to cellular control mGAPDH. 3 biological replicates, Wilcoxon Test. Normality determined by Kolmogorov-Smirnov test. *, P□<□0.05; **, P□<□0.01. ns, not significant.

We also produced lentiviruses expressing shRNAs targeting the core components of PRC1, Ring1A and Ring1B. Depletion was validated by both RT-qPCR and semi-quantitative immunofluorescence (Fig. 4E and 4F, and Supplemental Figure 1 G and 1H). Neonatal sympathetic neurons were transduced 5 days prior to infection with HSV-1 Stayput-GFP, and viral gene expression was analyzed at 3 days post-infection. The co-depletion of both Ring1A and Ring1B was also carried out because previous studies have shown redundancy between the two proteins in vPRC1 activity (41). For each class of viral lytic genes, viral gene expression was enhanced with the single knockdown of Ring1A (via two independent shRNA-targeting lentiviruses) and dual Ring1A/B knockdown (Fig 4D, Supplemental Figure 1 E and F). These functional data in combination with our kinetic data suggest that PRC1 and H2AK119ub are deposited onto the HSV-1 genome early during latency establishment to silence viral lytic genes.

We also depleted the core component of PRC2, Suz12, using one of three independent shRNA-targeting lentiviruses and subsequently infected cultures. It has previously been demonstrated that Suz12 is required for PRC2 activity and that its elimination prevents its function (42, 43). Upon Suz12 depletion, as validated at the level of RNA during each individual replicate in Fig 4G, the expression of all three classes of viral genes analyzed did not demonstrate a robust or significant enhancement at 3 days post-infection (Fig 4D, Supplemental Figure 1E and 1F). Therefore, our data suggest that, in contrast to PRC1, PRC2 does not promote HSV-1 lytic gene silencing during the establishment of a latent infection.

### PRC1 inhibition reduces future reactivation *in vitro*

Only a proportion of HSV-1 genomes associated with H2AK119ub, Ring1B, and H3K27me3 during established latency. We and others have also demonstrated that only a proportion of individual latent genomes initiate reactivation in response to different stimuli (21, 22, 30–32, 44). Our recent study found that genomes associated with H3K9me3 and read by ATRX are restricted for reactivation (26). Therefore, we hypothesized that association with H2Aub1 and/or H3K27me3 may result in genomes more primed for reactivation.

We therefore investigated the ability of viral genomes that undergo Polycomb silencing to reactivate by inhibiting PRC1 or PRC2 during latency establishment in the presence of viral DNA replication inhibitor acyclovir (ACV). The use of ACV here permits the equal establishment of latent infection despite PRC1/2 inhibitor treatment because viral DNA copy number remains stable (Fig. 5A). Following latency establishment in the presence of PRC1/2 inhibitors, cultures were stimulated with LY294002, and the number of GFP-positive neurons at 48 hours post-stimulus was quantified as a readout of neurons that have undergone the full HSV-1 reactivation. Interestingly, when PRC1 activity was inhibited during the establishment of latency, future reactivation was robustly decreased (Fig 5B). This phenotype was recapitulated using three independent PRC1 inhibitors. As this reduction in reactivation occurred despite equivalent latent viral DNA copy numbers, the reduced capacity to reactivate under PRC1 inhibition was not due to fewer viral genomes that established a latent infection. Notably, this reactivation defect was not observed when PRC2 was inhibited during this timeframe using three independent inhibitors. Further, reactivation was not perturbed when RB-3 was added instead for three days at 3-days post-infection (Fig. 5C), confirming that interrupting with Ring1 activity as the H2Aub1 modification is being depositing within the first 3 days of infection is required to impact later reactivation.

**Figure 5.**
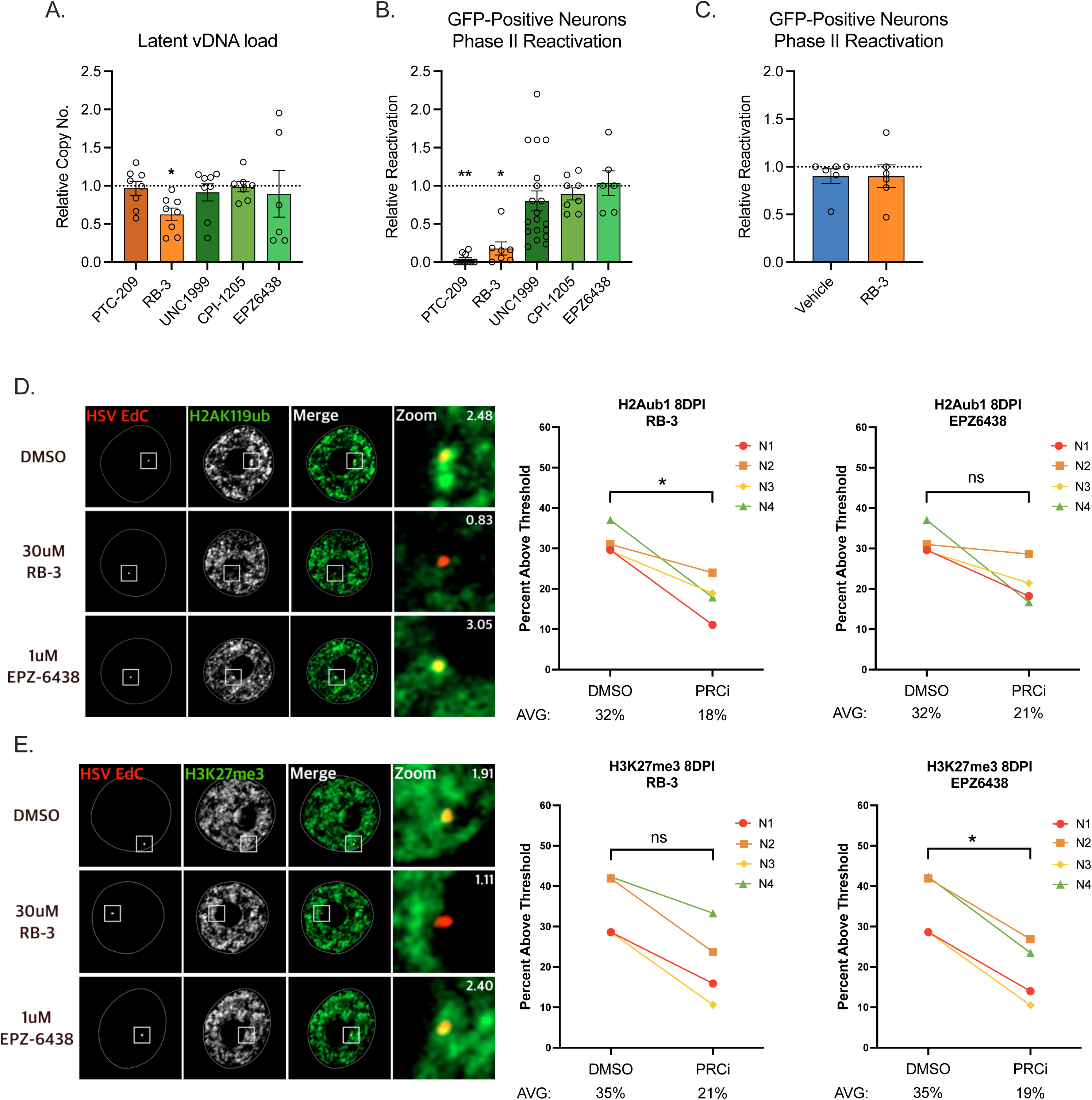
PRC1 activity during latency establishment is required to establish a reactivation-competent HSV-1 latent infection. Primary neurons were treated with a vehicle control, or with inhibitors of PRC1 (warm colors; PRT4165 40 μM, PTC-209 2 μM, RB-3 30 μM), or PRC2 (cool colors; UNC1999 1 μM, CPI-1205 10 μM, 1 μM EPZ6438) pre-infection for 1 hour, during inoculation with Stayput-GFP in the presence of ACV, and following the removal of inoculum. 6 days following infection, ACV and respective inhibitors were removed from cultures, and spontaneous reactivation, as indicated by GFP-positive neurons, was monitored for two days. Latently infected cultures were reactivated with LY294002 and GFP-positive neurons indicative of full reactivation events were monitored over time. (A) Latent viral DNA copy number normalized to vehicle controls. (B) The number of GFP-positive neurons at 48 hours post-stimulus presented as a fraction of vehicle control reactivation output. 3 biological replicates; Wilcoxon test against respective controls. Normality determined by Kolmogorov-Smirnov test. *, P□<□0.05; **, P□<□0.01. ns, not significant. D) Quantification of genome co-localization with H2Aub1 following RB-3 and EPZ6438 treatment E). Quantification of genome co-localization with H3K27me3 following RB-3 and EPZ6438 treatment.

We verified the loss of H2AK119ub or H3K27me3 by quantifying the genome co-localization in the presence of the PRC1 inhibitor RB-3 or the PRC2 inhibitor EPZ6438. We found that RB-3 significantly reduced the proportion of genomes associated with H2Aub1 from 32% to 18% positive co-localization, a value similar to that expected by chance (Fig. 5D). H3K27me3 co-localization was reduced by EPZ-6838 treatment, from 35% to 19% (Fig. 5E). We also investigated the effects of PRC1 inhibition on H3K27me3 levels and of PRC2 inhibition on H2Aub1 levels. RB-3 treatment resulted in a small and consistent decrease in H3K27me3 levels, from 35 to 21%. However, this decrease was not significant (Fig. 5E). This indicates that on the viral genome, either H2Aub1 deposition is required for H3K27me3 deposition on a subpopulation of neurons, or that H3K27me3 deposition is independent of vPRC activity. Inhibition of PRC2 also had a small but non-significant effect on H2Aub1 co-localization, decreasing from 32% to 21%. However, this was not fully convincing between individual experiments (Fig. 5D). Therefore, based on this data alone, it is hard to conclude for sure the impact of PRC2 activity on H2Aub1. Importantly, our co-localization analysis verifies both the reduced association of h2Aub1 with the genome in the presence of the PRC1 inhibitor and demonstrates that Polycomb silencing, specifically H2AK119ub deposition, during the early phase of latency establishment promotes later reactivation, possibly via the formation of more reactivation-competent latent HSV-1 genomes.

## Discussion

Here, we demonstrate that PRC1-mediated H2AK119ub is enriched on the HSV-1 genome during latency established *in vivo* and *in vitro*. The direct contribution of heterochromatin deposition on the viral genome to promoting gene repression and establishment of latency has not been described. Here we dissect the contributions of PRC1 and PRC2 and find that only the loss of inhibition affects viral gene expression, and only mildly. These data suggest that H2Aub1 only moderately represses gene expression, or, following PRC1 inhibition, alternative forms of heterochromatin can participate in gene silencing instead. This latter explanation is most intriguing, as we found that although PRC1 inhibition had only mild effects on latency establishment, its inhibition had a drastic effect on the ability of genomes to reactivate. Therefore, the type of heterochromatin on the viral genome may play a less important role in promoting gene silencing but does impact the ability of viral genomes to reactivate.

Interestingly, the enrichment of PRC1 component BMI1 was previously explored on the latent HSV-1 genome in a mouse model, but not found to be significant (2, 4). However, it has become clear over time from studies on the cellular genome that Polycomb repressor complexes can be found in a multitude of forms (45–47). BMI1 traditionally associates with one form of PRC1 (cPRC1) that does not efficiently carry out H2AK119ub (13, 14, 48, 49). Therefore, the enrichment of H2AK119ub but not BMI on the HSV-1 genome is not confounding. What is unexpected is that H2AK119ub is actively targeted to the HSV-1 genome in the specialized cell type of a post-mitotic peripheral neuron. Studies on the cellular genome have thus far suggested that vPRC1-mediated H2AK119ub deposition and silencing is abundant in embryonic cells, like neuronal progenitor cells/NPCs, but decreases in relevance to cellular genome silencing and maintenance as cells differentiate (15). Our work now demonstrates that in response to environmental stimuli, such as a viral infection, H2AK119ub deposition is a relevant process in post-mitotic neurons. We also confirm the kinetics of H3K27me3 enrichment previously observed *in vivo*, where the HSV-1 genome becomes enriched in H3K27me3 at later time points post-infection after lytic genes have been repressed (2, 4).

These are also the first functional studies that directly probe the contributions of PRC1/2 to lytic viral gene silencing during latency establishment. Importantly, we find that H2AK119ub is targeted to the viral genome and contributes to the repression of lytic genes during the early stages of latency establishment. Consistent with its kinetics, PRC2 does not actively repress viral gene expression, at least not during the time points tested here. These data, taken together, suggest a model in which PRC1 is targeted to the viral genome to initiate viral gene repression. H3K27me3, either dependent on PRC1 or via PRC1-independent mechanisms, is subsequently recruited to reinforce and maintain viral gene repression. On the cellular genome, PRC2 can be recruited in a manner that is dependent on prior H2AK119ub deposition, or independently of H2AK119ub (50–52). This can result in different types of Polycomb silencing that can be either enriched or depleted for H2AK119ub. We observed a similar proportion of co-localization between H2AK119ub and H3K27me3 and the latent genome (one-third), suggesting that these modifications bind to the same viral genomes; however, we cannot currently rule out that they are on separate genomes. The role of H2AK119ub in promoting H3K27me3 has predominately been studied in embryonic stem cells and during the early stages of differentiation (51–55). The protein that promotes PRC2 recruitment to H2AK119ub to mediate H3K27me3 addition (JARID2) is altered in differentiated cells in that a truncated version predominates that lacks both the H3AK119ub reader domain and PRC2 binding domain (55, 56). Therefore, it remains unresolved whether H2AK119ub1 can directly promote H3K27me3 in neurons and on the HSV genome, and the contribution of JARID2.

The mechanisms of PRC1, as well as PRC2 recruitment to the HSV-1 genome remain unresolved. One potential mechanism of recruitment and/or activation of PRC1 and PRC2 to mediate deposition of histone post-translational modifications is via binding to long noncoding RNAs (lncRNAs). Notably, during the establishment and maintenance of HSV-1 latent infection, the virus expresses a lncRNA LAT, at least in a proportion of neurons. The exact molecular functions of the LAT are unknown, and while it is not essential for the establishment of a latent infection, it may modulate lytic gene expression and promote increased association with H3K27me3 (3). Viral LAT expression may therefore actively promote a type of gene silencing that is capable of reactivating. In fact, the LAT has been demonstrated to enhance reactivation in several models, further supporting this theory (57, 58). PRC1 may alternatively be recruited using host protein KDM2B, which recruits vPRC1 to cellular genomes by binding unmethylated GC-rich sequences (13, 59), which comprise much of the HSV-1 and other human herpesvirus genomes (60). In fact, fellow human herpesvirus KSHV uses its intrinsic GC-richness and KDM2B to recruit Polycomb (61) (62). PRC1 may also be recruited to the HSV-1 genome by host transcription factors, such as RUNX1, which is abundantly expressed in in sensory neurons and sympathetic neurons, where HSV-1 readily establishes latency. RE1-silencing transcription factor (REST, also known as neuron-restrictive silencing factor (NRSF)) can also interact with PRC1 (and PRC2) and maintains Polycomb binding to neuronal genes in the human teratocarcinoma NT2-D1 cells and mESCs (63). The HSV-1 genome carries predicted REST binding sites and may repress the ability of HSV-1 to transcribe lytic viral genes and reactivate (64) (65).

PRC1-mediated silencing may be more amenable to reactivation, but the question of how repressive H2AK119ub is modified or removed following a reactivation stimulus remains unresolved. Accumulating evidence increasingly suggests that HSV-1 reactivation is a biphasic process. Initial viral gene expression events during Phase I are independent of the removal of repressive heterochromatin, which is instead modified to enable viral gene expression. Repressive H3K9me3, for example, undergoes a phospho-methyl switch, where histone phosphorylation on a neighboring serine allows viral gene expression to occur despite the persistence of repressive H3K9 (66). During Phase II, repressive heterochromatin is fully removed. One model is that during Phase II reactivation, H2AK119ub would likely be fully removed from the viral genome by de-ubiquitinase (DUB) BRCA1-associated protein 1, BAP1. Interestingly, BAP1 has previously been demonstrated to associate with host cell factor 1 (HCF-1), which may also be required for the progression to the full, Phase II reactivation by interacting with viral protein VP16 (67–70).

## Materials & Methods

### Reagents

Compounds used in the study are described in Table 1. Compound concentrations were used based on previously published IC50s and assessed for neuronal toxicity using the cell body and axon health and degeneration index as previously published (30). All compounds had an average score less than or equal to 1.

**Table 1.**
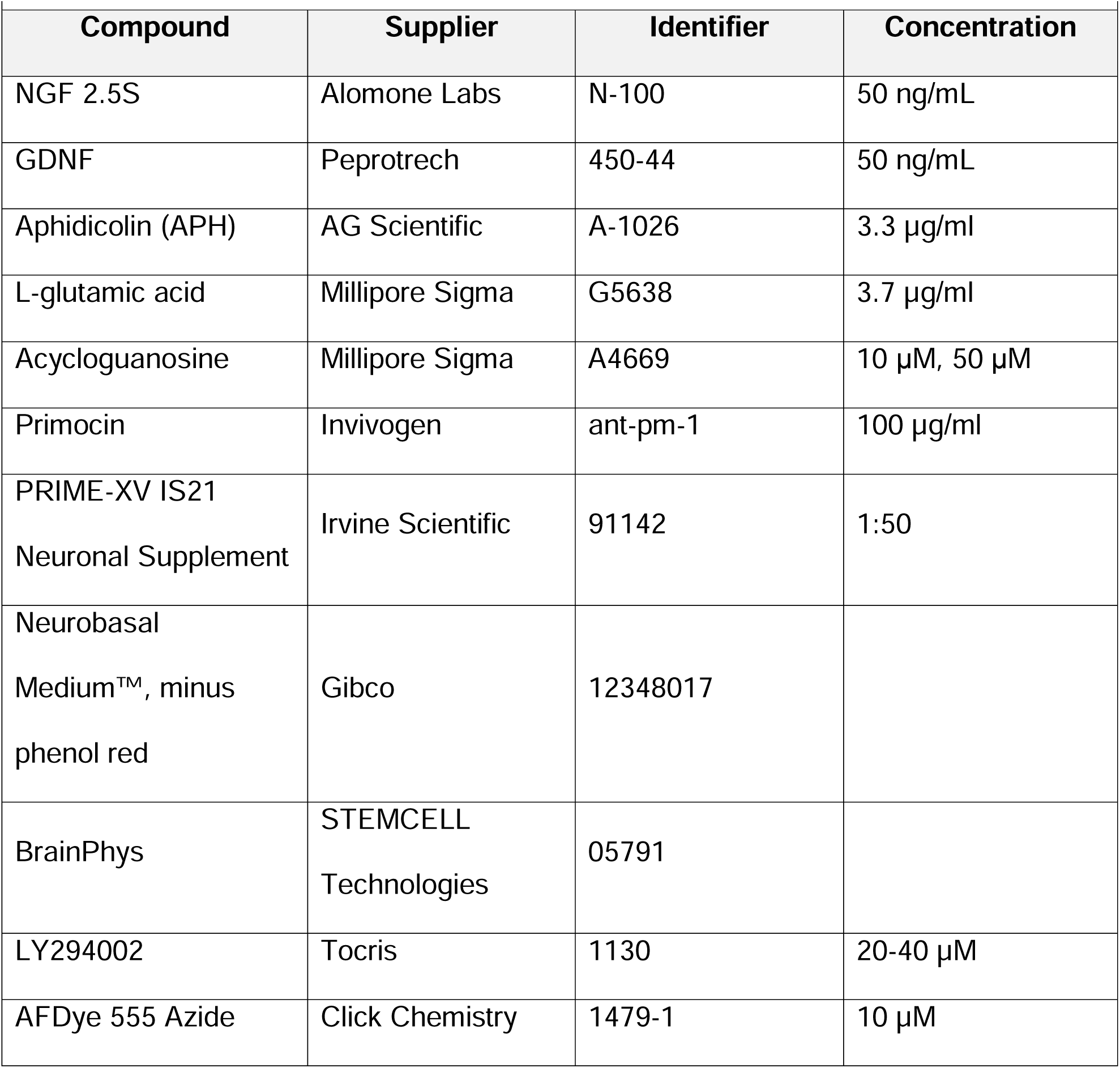
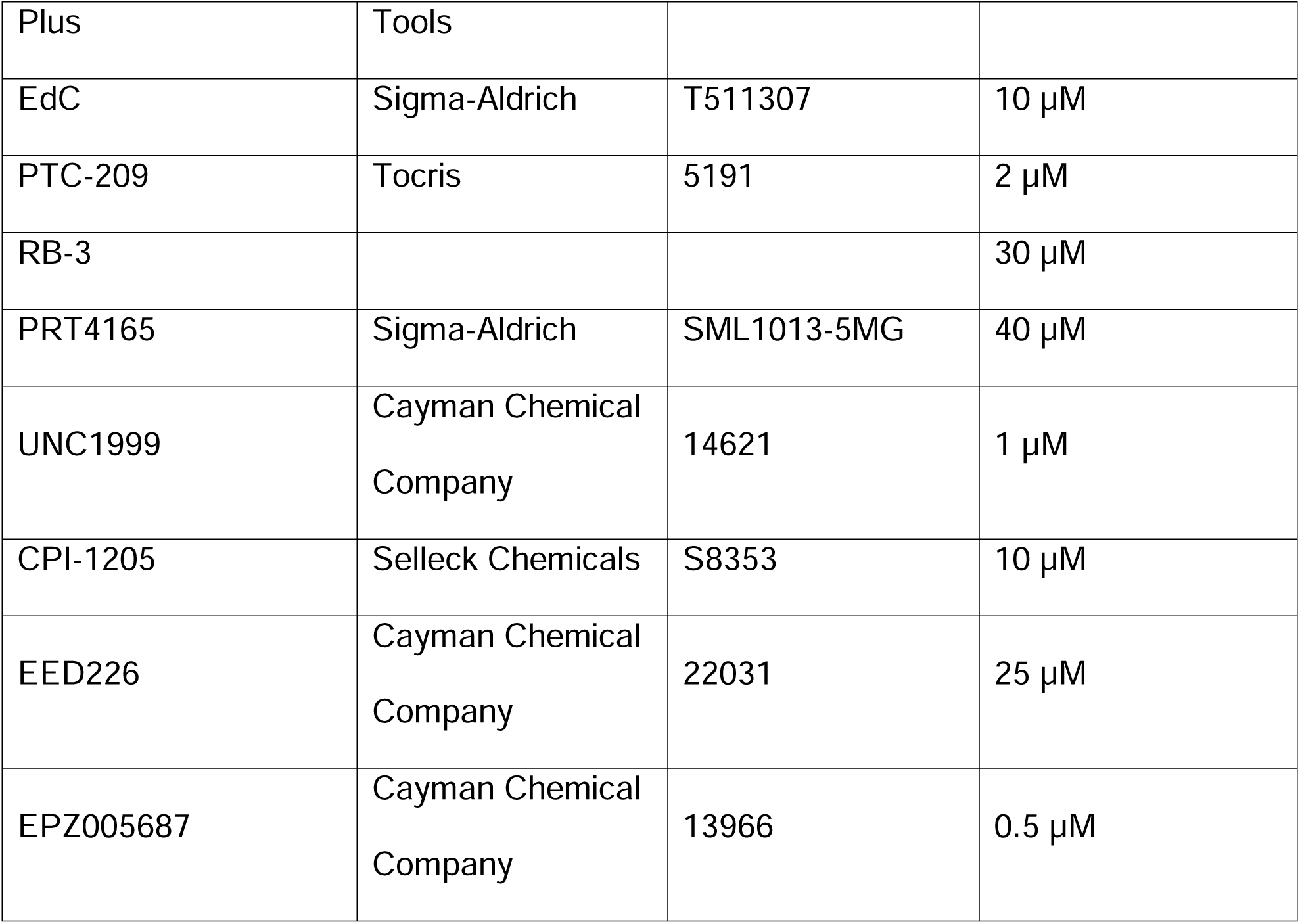
Compounds used and concentrations.

### Preparation of HSV-1 virus stocks

Stayput-GFP, as well as SCgHZ, is propagated and titrated on previously constructed gH-complementing F6 cells, which contain copies of the gH gene under the control of an HSV-1 gD promoter, as described in (71). Vero F6 cells were maintained in Dulbecco’s Modified Eagle’s Medium (Gibco) supplemented with 10% FetaPlex (Gemini BioProducts) and 250 μg/mL of G418/Geneticin (Gibco).

HSV-1 stocks of Syn17+, and KOS (for *in vivo* experiments) were grown and titrated on Vero cells obtained from the American Type Culture Collection (Manassas, VA). Cells were maintained in Dulbecco’s Modified Eagle’s Medium (Gibco) supplemented with 10% FetalPlex (Gemini Bio-Products) and 2 mM L-Glutamine. eGFP-Us11 Patton (HSV-1 Patton strain with eGFP reporter protein fused to true late protein Us11 (72) was kindly provided by Dr. Ian Mohr at New York University.

### Primary neuronal cultures

Sympathetic neurons from the superior cervical ganglia (SCG) of post-natal day 0–2 (P0-P2) CD1 Mice (Charles River Laboratories) were dissected as previously described (66). Sensory neurons from the trigeminal ganglia (TG) or adult (P21-24) CD1 Mice (Charles River Laboratories) were dissected as previously described (73). Rodent handling and husbandry were carried out under animal protocols approved by the Animal Care and Use Committee of the University of Virginia (UVA). Ganglia were briefly kept in Leibovitz’s L-15 media with 2.05 mM L-Glutamine before dissociation in Collagenase Type IV (1 mg/mL) followed by Trypsin (2.5 mg/mL) for 20 min each at 37 C. Dissociated ganglia were triturated, and approximately 5-10,000 neurons per well were plated onto rat tail collagen in a 24-well plate. Sympathetic neurons were maintained in CM1 (Neurobasal Medium supplemented with PRIME-XV IS21 Neuronal Supplement (Irvine Scientific), 50 ng/mL Mouse NGF 2.5S, 2 mM L-Glutamine, and Primocin). Aphidicolin (3.3 μg/mL) was added to the CM1 for the first 5 days post-dissection to select against proliferating cells. Sensory neurons were maintained in the same media supplemented with GDNF (50 ng/ml; Peprotech 450-44), and more Aphidicolin (6.6 μg/mL) was used for the first 5 days as more non-neuronal cells tend to be dissected with this neuron type.

### Establishment and reactivation of latent HSV-1 infection in primary neurons

P6-8 SCG neurons were infected with Stayput Us11-GFP at MOI 7.5 PFU/cell, assuming 10,000 cells per well in Dulbecco’s Phosphate Buffered Saline (DPBS) + CaCl_2_ + MgCl_2_ supplemented with 1% fetal bovine serum and 4.5 g/L glucose for 3.5 h at 37°C. Post infection, inoculum was replaced with feeding media (as described above). Acyclovir (ACV) was not utilized in these infections. Lytic infection was quantified by performing reverse transcription–quantitative PCR (RT–qPCR) of HSV-1 lytic mRNAs isolated from the cells in culture.

Neonatal SCGs were infected at postnatal days 6-8 with either Us11-GFP, SCgHZ, or Stayput-GFP at an MOI of 7.5 PFU/cell assuming 10,000 cells/well in DPBS +CaCl_2_+MgCl_2_ supplemented with 1% Fetal Bovine Serum, 4.5 g/L glucose, and either with or without 10 μM Acyclovir (ACV) for 3 hr at 37 C. Post infection, inoculum was replaced with feeding media with or without 50 μM ACV. Reactivation was carried out in BrainPhys (Stem Cell Technologies) supplemented with 2 mM L-Glutamine, 10% Fetal Bovine Serum, Mouse NGF 2.5S (50 ng/mL) and Primocin. Reactivation was quantified by counting number of GFP-positive neurons or performing Reverse Transcription Quantitative PCR (RT-qPCR) for HSV-1 lytic transcripts.

### Mouse Infections

Five-week-old male and female CD-1 mice (Charles River Laboratories) were anesthetized by intraperitoneal injection of ketamine hydrochloride (80 mg/kg of body weight) and xylazine hydrochloride (10 mg/kg) and inoculated with 1.5 × 10^6^ PFU of HSV-1 WT KOS/eye of virus (in a 5 μL volume) onto scarified corneas, as described previously (2). Mice were housed in accordance with institutional and National Institutes of Health guidelines on the care and use of animals in research, and all procedures were approved by the Institutional Animal Care and Use Committee of the University of Virginia. Criteria used for clinical scoring based on the formation of lesions and neurological and eye symptoms are shown in Table 2 and were based on a previously established scoring scale (74). Mice were randomly assigned to groups, and all experiments included biological replicates from independent litters.

**Table 2.**
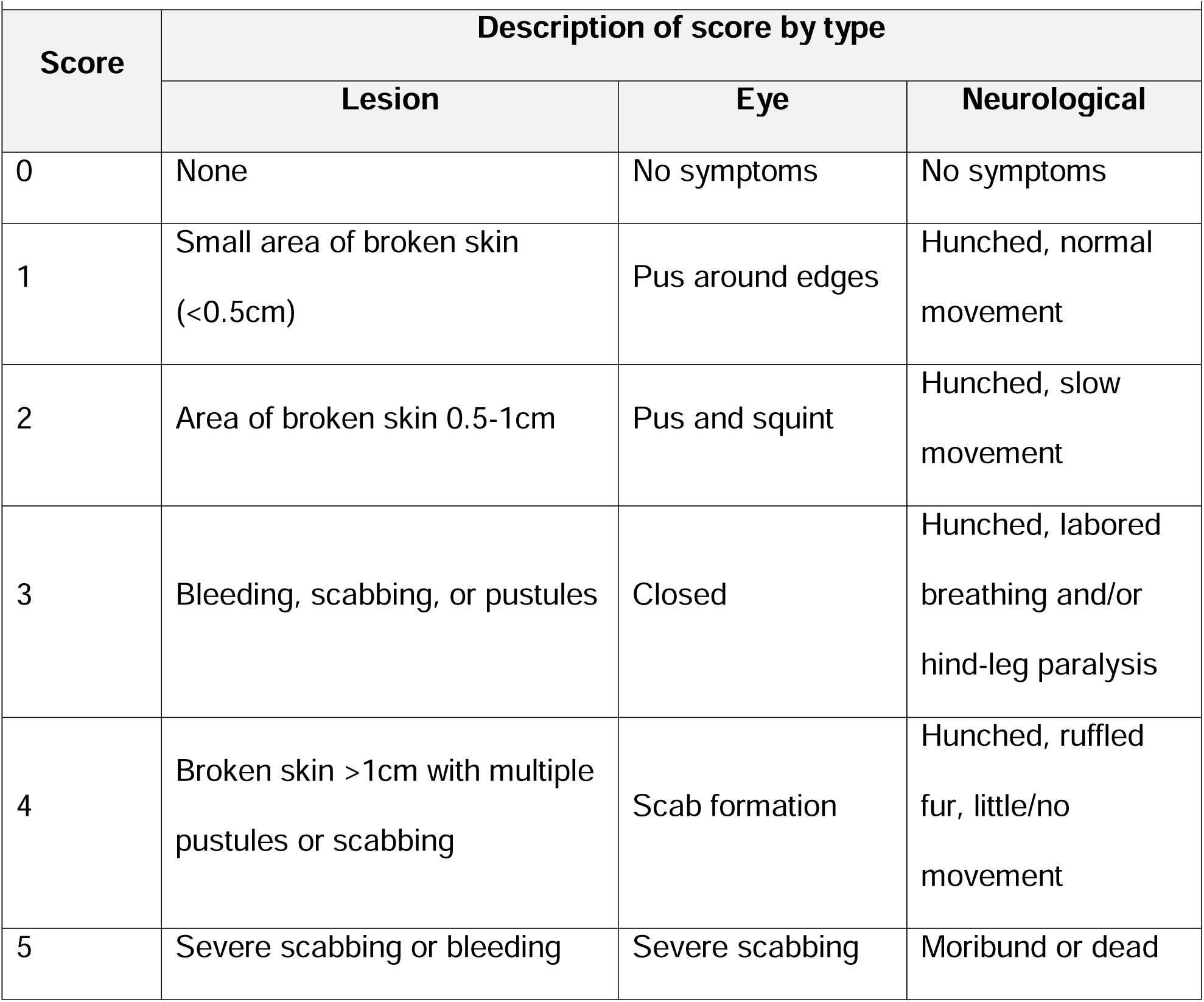
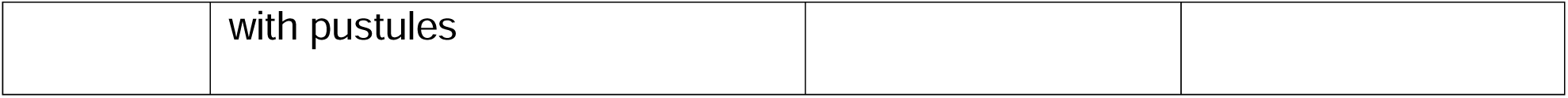
Clinical scoring scale for HSV-infected mice.

**Table 3.** Antibodies.

| **Table 3. Antibodies** |  |  |  |
| --- | --- | --- | --- |
| **Designation** | **Source or reference** | **Identifiers** | **Additional Information** |
| Anti-α-tubulin (Mouse monoclonal) | Millipore sigma | Cat #T9026 | WB (1:2500) |
| Anti-Rabbit IgG Antibody (H+L), Peroxidase (Goat polyclonal) | Vector Labs | Cat #PI-1000 | WB (1:10,000) |
| Anti-mouse IgG Antibody (H+L), Peroxidase (Horse polyclonal) | Vector Labs | Cat #PI-2000 | WB (1:10,000) |
| Beta III Tubulin | EMD Millipore | Cat #9354 | IF (1:500) |
| IgG | EMD Millipore | 12-370 | CUT&RUN<br>ChIP-qPCR |
| H2A | Cell Signaling Technology | 12349S | ChIP-qPCR |
| H2AK119ub | Cell Signaling Technology | 8240S | ChIP (1:100)<br>CUT&RUN<br>IF (1:1600) |
| H3K27me3 | Cell Signaling Technology | Cat #9733 | CUT&RUN<br>ChIP 1:50<br>IF (1:750) |
| H3K27me2 | Active Motif | Cat #39245 | IF 1:500 |
| Ring1B | Cell Signaling Technology | 5694S | IF, WB |
| ICP4 | Abcam | ab6514 | IF (1:850) |
| ICP5 | Abcam | ab6508 | IF |
| ICP8 | Abcam | ab20194 | IF |
| H3 | Cell Signaling Technology |  | WB |
| Alpha-tubulin | Cell Signaling Technology |  | WB |

### Western Blotting

Neurons were lysed in RIPA Buffer with cOmplete, Mini, EDTA-Free Protease Inhibitor Cocktail (Roche) and PhosSTOP Phosphatase Inhibitor Cocktail (Roche) on ice for 2 hours with regular vortexing to aid lysis. Insoluble proteins were removed via centrifugation, and lysate protein concentration was determined using the Pierce Bicinchoninic Acid Protein Assay Kit (Invitrogen) using a standard curve created with BSA standards of known concentration. Equal quantities of protein (15–50 μg) were resolved on 4–20% gradient SDS-Polyacrylamide gels (Bio-Rad) and then transferred onto Polyvinylidene difluoride membranes (Millipore Sigma). Membranes were blocked in PVDF Blocking Reagent for Can Get Signal (Toyobo) for 1 hr. Primary antibodies were diluted in Can Get Signal Immunoreaction Enhancer Solution 1 (Toyobo) and membranes were incubated overnight at 4°C. HRP-labeled secondary antibodies were diluted in Can Get Signal Immunoreaction Enhancer Solution 2 (Toyobo) and membranes were incubated for 1 hr at room temperature. Antibody usage is recorded in Table 4. Blots were developed using Western Lightning Plus-ECL Enhanced Chemiluminescence Substrate (PerkinElmer) and ProSignal ECL Blotting Film (Prometheus Protein Biology Products) according to manufacturer’s instructions. Blots were stripped for reblotting using NewBlot PVDF Stripping Buffer (Licor). Band density was quantified in ImageJ.

**Table 4.**
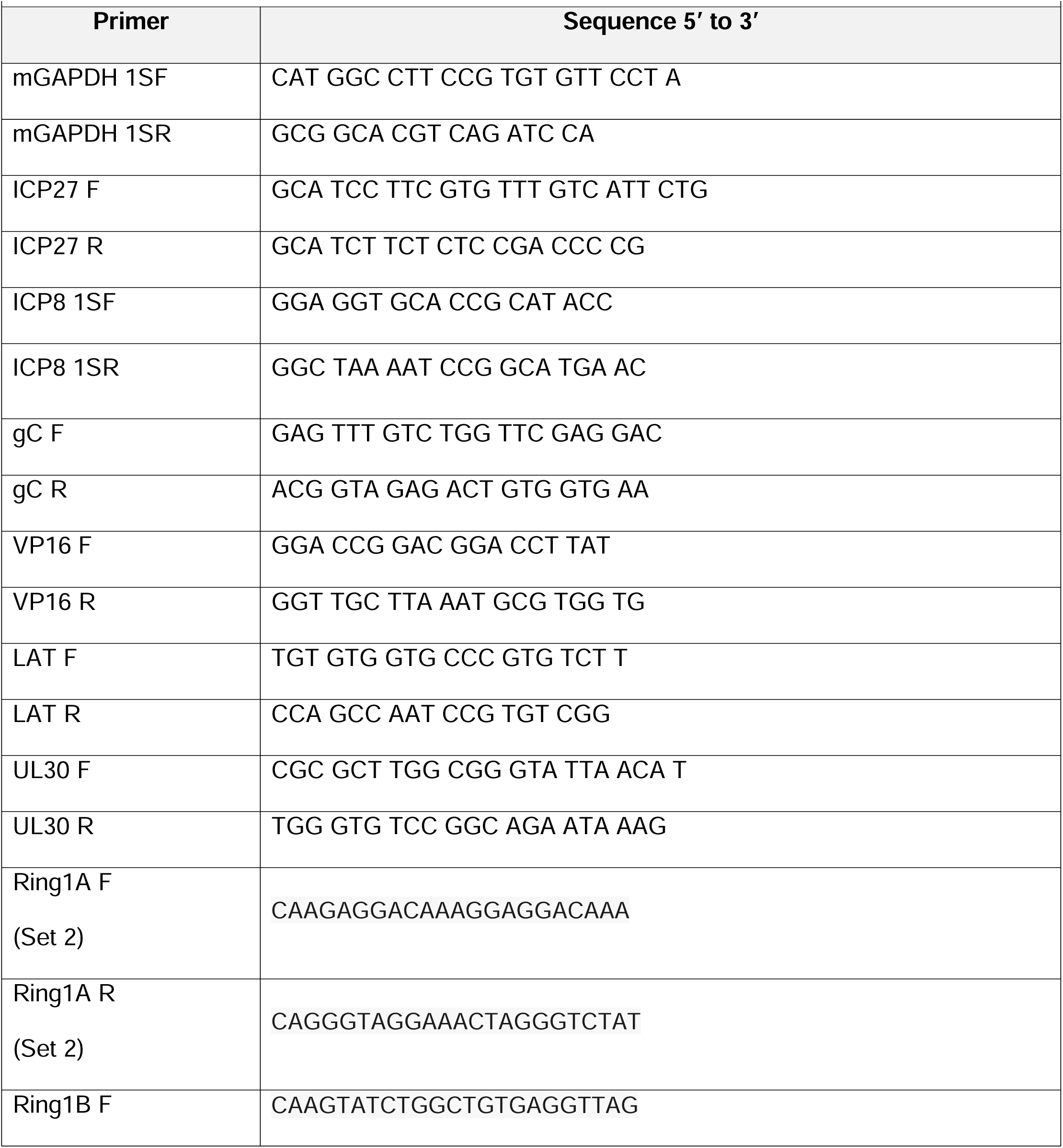
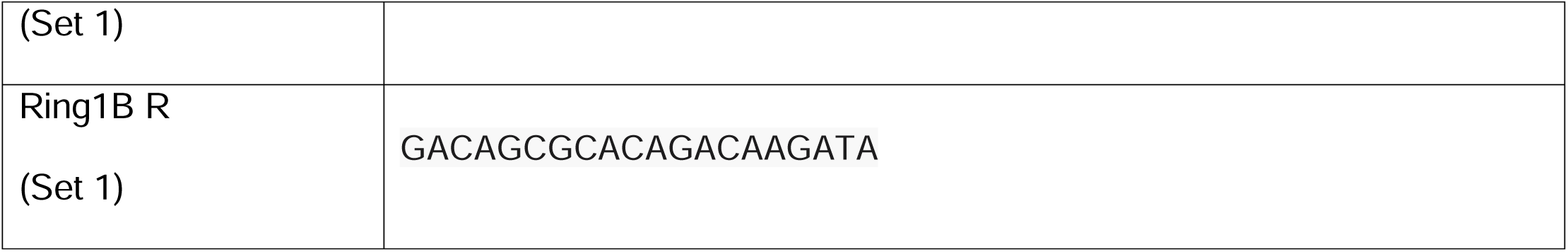
Sequences of Primers Used.

### Preparation of lentiviral vectors

Lentiviruses expressing shRNA against Ring1A (Ring1A-1 = TRCN0000040561 Ring1A-2 = TRCN0000040562), Ring1B (Ring1B-1 = TRCN0000226018, Ring1B-2= TRCN0000226019), SUZ12 (Suz12-1 = TRCN0000234040, Suz12-2 = TRCN0000218848, Suz12-3 =TRCN0000123893) or a control lentivirus shRNA (empty pLKO.1 or pLKO.5 vector) were prepared by co-transfection with psPAX2 and pCMV-VSV-G (75) using the 293LTV packaging cell line (Cell Biolabs). Supernatant was harvested at 40- and 64-h post-transfection and filtered using a 45 μM PES filter. Sympathetic neurons were transduced overnight in neuronal media containing 8 μg/ml protamine sulfate and 50 μM ACV.

### CUT&RUN Sequencing

A minimum of approximately 120,000 neurons were plated for each CUT&RUN reaction. Neurons were collected following 10-minute incubation with trypsin at 37 C without fixation. Neurons were permeabilized with 0.01% digitonin (concentration optimized for neurons) and CUT&RUN was performed using reagents included in the CUT&RUN Kit (EpiCypher) according to attached protocol. Periodic quality checks as recommended and detailed by protocol Appendix were performed and passed for neurons. CUT&RUN libraries were prepared using CUT&RUN Library Prep Kit (EpiCypher) and subject to pair-ended partial-lane sequencing and de-multiplexing using NovaSeq (Novogene). Data analysis was performed using command line and R code, and workflow, adapted from tutorial on IO protocols (Zheng Y et al (2020) Protocol.io). The Rivanna high-performance computing environment (UVA Research Computing) was used for command line data processing. The HSV-1 SC16 genome sequence (NCBI NC_KX946970.1) was used in combination with the mm39 mouse genome assembly (RefSeq NC_000071.7) to make a joint Bowtie2 index genome.

Sequence alignment was performed with bowtie2 with the following settings*: --end-to-end --very-sensitive --no-mixed --no-discordant --phred33 -I 10 -X 700.* Separate alignments was performed to spike-in *E. Coli* DNA (MG1655, Genbank U00096.3), from which sequencing depth was calculated and reads normalized accordingly before filtering into separate human and viral bedgaph files. Data quality control and visualization were performed using R. Two experimental replicates were performed.

### Click Chemistry

For EdC-labeled HSV virus infections, an MOI of 7.5 was used. EdC labelled virus was prepared using a previously described method (76). Click chemistry was carried out as described previously (77) with some modifications. Neurons were washed with CSK buffer (10 mM HEPES, 100 mM NaCl, 300 mM Sucrose, 3 mM MgCl_2_, 5 mM EGTA) and simultaneously fixed and permeabilized for 10 minutes in 1.8% methanol-free formaldehyde (0.5% Triton X-100, 1% phenylmethylsulfonyl fluoride (PMSF)) in CSK buffer, then washed twice with PBS before continuing to the click chemistry reaction and immunostaining. Samples were blocked with 3% BSA for 30 minutes, followed by click chemistry using EdC-labelled HSV DNA and the Click-iT EdU Alexa Flour 555 Imaging Kit (ThermoFisher Scientific, C10638) according to the manufacturer’s instructions. For immunostaining, samples were incubated overnight with primary antibodies in 3% BSA.

Following primary antibody treatment, neurons were incubated for one hour in Alexa Fluor® 488-, 555-, and 647-conjugated secondary antibodies for multi-color imaging (Invitrogen). Nuclei were stained with Hoechst 33258 (Life Technologies). Images were acquired at 60x using an sCMOS charge-coupled device camera (pco.edge) mounted on a Nikon Eclipse Ti Inverted Epifluorescent microscope using NIS-Elements software (Nikon). Images were analyzed and intensity quantified using ImageJ.

### Immunofluorescence

Neurons were fixed for 15 min in 4% Formaldehyde and blocked in 5% Bovine Serum Albumin and 0.3% Triton X-100 and incubated overnight in primary antibody. Following primary antibody treatment, neurons were incubated for 1 hr in Alexa Fluor 488-, 555-, and 647-conjugated secondary antibodies for multi-color imaging (Invitrogen). Nuclei were stained with Hoechst 33258 (Life Technologies). Images were acquired using an sCMOS charge-coupled device camera (pco.edge) mounted on a Nikon Eclipse Ti Inverted Epifluorescent microscope using NIS-Elements software (Nikon). Images were analyzed using ImageJ.

### Analysis of mRNA expression and viral genome copy number

To assess expression of HSV-1 lytic mRNA, total RNA was extracted from approximately 5,000 neurons using the Quick-RNA Miniprep Kit (Zymo Research) with an on-column DNase I digestion. mRNA was converted to cDNA using the Maxima cDNA synthesis kit (Thermo Fisher) using random hexamers for first-strand synthesis and equal amounts of RNA (20–30 ng/ reaction). To assess viral DNA load, total DNA was extracted from approximately 5,000 neurons using the Quick-DNA Miniprep Plus Kit (Zymo Research). qPCR was carried out using PowerUp^TM^ SYBR Green Master Mix (Thermoscientific). Relative mRNA/DNA copy numbers were determined using the Comparative CT (ΔΔCT) method normalized to mRNA levels in latently infected samples. Viral RNAs were normalized to mouse reference gene 18s ribosomal RNA. All samples were run in duplicate on an Applied Biosystems QuantStudio 6 Flex Real-Time PCR System and the mean fold change compared to the reference gene calculated.

Exact copy numbers were determined by comparison to standard curves of known DNA copy number of viral genomes or plasmid containing 18S rRNA (p18S.1 tag was a gift from Vincent Mauro; Addgene plasmid # 51729 ; http://n2t.net/addgene:51729 ; RRID:Addgene_51729, (78).

### Statistical Analysis

All statistical analyses were performed using Prism V10. The normality test and statistical test used for each figure are denoted in the figure legends. Individual biological replicates are plotted, each corresponding to a single well of cells/neurons. All neuronal experiments were repeated from pooled neurons from at least 3 litters.

## References

1. R. F. Itzhaki, Herpes simplex virus type 1 and Alzheimer’s disease: increasing evidence for a major role of the virus. Front Aging Neurosci 6, 202 (2014).

2. A. R. Cliffe, D. M. Coen, D. M. Knipe, Kinetics of facultative heterochromatin and polycomb group protein association with the herpes simplex viral genome during establishment of latent infection. mBio 4 (2013).

3. A. R. Cliffe, D. A. Garber, D. M. Knipe, Transcription of the herpes simplex virus latency-associated transcript promotes the formation of facultative heterochromatin on lytic promoters. J Virol 83, 8182–8190 (2009).

4. D. L. Kwiatkowski, H. W. Thompson, D. C. Bloom, The polycomb group protein Bmi1 binds to the herpes simplex virus 1 latent genome and maintains repressive histone marks during latency. J Virol 83, 8173–8181 (2009).

5. M. P. Nicoll, et al., The HSV-1 Latency-Associated Transcript Functions to Repress Latent Phase Lytic Gene Expression and Suppress Virus Reactivation from Latently Infected Neurons. PLoS Pathog 12, e1005539 (2016).

6. R. Cao et al., Role of histone H3 lysine 27 methylation in Polycomb-group silencing. Science 298, 1039–1043 (2002).

7. P. Trojer, D. Reinberg, Facultative heterochromatin: is there a distinctive molecular signature? Mol Cell 28, 1–13 (2007).

8. R. Cao, Y. Tsukada, Y. Zhang, Role of Bmi-1 and Ring1A in H2A ubiquitylation and Hox gene silencing. Mol Cell 20, 845–854 (2005).

9. M. de Napoles et al., Polycomb group proteins Ring1A/B link ubiquitylation of histone H2A to heritable gene silencing and X inactivation. Dev Cell 7, 663–676 (2004).

10. H. Wang et al., Role of histone H2A ubiquitination in Polycomb silencing. Nature 431, 873–878 (2004).

11. N. P. Blackledge et al., PRC1 Catalytic Activity Is Central to Polycomb System Function. Mol Cell 77, 857–874.e859 (2020).

12. S. Tamburri et al., Histone H2AK119 Mono-Ubiquitination Is Essential for Polycomb-Mediated Transcriptional Repression. Mol Cell 77, 840–856 e845 (2020).

13. N. P. Blackledge et al., Variant PRC1 complex-dependent H2A ubiquitylation drives PRC2 recruitment and polycomb domain formation. Cell 157, 1445–1459 (2014).

14. L. Tavares et al., RYBP-PRC1 complexes mediate H2A ubiquitylation at polycomb target sites independently of PRC2 and H3K27me3. Cell 148, 664–678 (2012).

15. M. Tsuboi et al., Ubiquitination-Independent Repression of PRC1 Targets during Neuronal Fate Restriction in the Developing Mouse Neocortex. Dev Cell 47, 758–772 e755 (2018).

16. N. P. Blackledge, R. J. Klose, The molecular principles of gene regulation by Polycomb repressive complexes. Nat Rev Mol Cell Biol 22, 815–833 (2021).

17. J. J. Kim, R. E. Kingston, Context-specific Polycomb mechanisms in development. Nat Rev Genet 10.1038/s41576-022-00499-0 (2022).

18. A. F. Lentine, S. L. Bachenheimer, Intracellular organization of herpes simplex virus type 1 DNA assayed by staphylococcal nuclease sensitivity. Virus Res 16, 275–292 (1990).

19. P. F. Pignatti, E. Cassai, Analysis of herpes simplex virus nucleoprotein complexes extracted from infected cells. J Virol 36, 816–828 (1980).

20. F. Catez et al., HSV-1 genome subnuclear positioning and associations with host-cell PML-NBs and centromeres regulate LAT locus transcription during latency in neurons. PLoS Pathog 8, e1002852 (2012).

21. S. Dochnal et al., DLK-Dependent Biphasic Reactivation of Herpes Simplex Virus Latency Established in the Absence of Antivirals. J Virol 96, e0050822 (2022).

22. J. B. Suzich et al., PML-NB-dependent type I interferon memory results in a restricted form of HSV latency. EMBO Rep 22, e52547 (2021).

23. N. M. Sawtell, R. L. Thompson, Comparison of herpes simplex virus reactivation in ganglia in vivo and in explants demonstrates quantitative and qualitative differences. J Virol 78, 7784–7794 (2004).

24. S. R. Cuddy et al., Neuronal hyperexcitability is a DLK-dependent trigger of herpes simplex virus reactivation that can be induced by IL-1. Elife 9 (2020).

25. J. Y. Kim, A. Mandarino, M. V. Chao, I. Mohr, A. C. Wilson, Transient reversal of episome silencing precedes VP16-dependent transcription during reactivation of latent HSV-1 in neurons. PLoS Pathog 8, e1002540 (2012).

26. A. L. Whitford et al., Interferon Dependent Immune Memory during HSV-1 Neuronal Latency Results in Increased H3K9me3 and Restriction of Reactivation by ATRX. bioRxiv 10.1101/2025.01.28.633013, 2025.2001.2028.633013 (2025).

27. S. A. Dochnal, A. K. Francois, A. R. Cliffe, De Novo Polycomb Recruitment: Lessons from Latent Herpesviruses. Viruses 13 (2021).

28. L. Condemi, I. Mocavini, S. Aranda, L. Di Croce, Polycomb function in early mouse development. Cell Death Differ 32, 90–99 (2025).

29. J. T. Proença, H. M. Coleman, V. Connor, D. J. Winton, S. Efstathiou, A historical analysis of herpes simplex virus promoter activation in vivo reveals distinct populations of latently infected neurones. J Gen Virol 89, 2965–2974 (2008).

30. S. R. Cuddy et al., Neuronal hyperexcitability is a DLK-dependent trigger of herpes simplex virus reactivation that can be induced by IL-1. Elife 9, e58037 (2020).

31. H. L. Hu et al., Single-cell transcriptomics identifies Gadd45b as a regulator of herpesvirus-reactivating neurons. EMBO Rep 23, e53543 (2022).

32. R. L. Thompson, C. M. Preston, N. M. Sawtell, De novo synthesis of VP16 coordinates the exit from HSV latency in vivo. PLoS Pathog 5, e1000352 (2009).

33. A. K. Francois et al., Single-genome analysis reveals a heterogeneous association of the herpes simplex virus genome with H3K27me2 and the reader PHF20L1 following infection of human fibroblasts. mBio 15, e0327823 (2024).

34. W. Zhou et al., Histone H2A monoubiquitination represses transcription by inhibiting RNA polymerase II transcriptional elongation. Mol Cell 29, 69–80 (2008).

35. J. K. Stock et al., Ring1-mediated ubiquitination of H2A restrains poised RNA polymerase II at bivalent genes in mouse ES cells. Nat Cell Biol 9, 1428–1435 (2007).

36. L. Di Croce, K. Helin, Transcriptional regulation by Polycomb group proteins. Nat Struct Mol Biol 20, 1147–1155 (2013).

37. N. P. Blackledge et al., PRC1 Catalytic Activity Is Central to Polycomb System Function. Mol Cell 77, 857–874 e859 (2020).

38. Y. Hou et al., PHF20L1 as a H3K27me2 reader coordinates with transcriptional repressors to promote breast tumorigenesis. Sci Adv 6, eaaz0356 (2020).

39. A. Kreso et al., Self-renewal as a therapeutic target in human colorectal cancer. Nat Med 20, 29–36 (2014).

40. S. Shukla et al., Small-molecule inhibitors targeting Polycomb repressive complex 1 RING domain. Nat Chem Biol 17, 784–793 (2021).

41. Y. Zhu et al., Functional redundancy among Polycomb complexes in maintaining the pluripotent state of embryonic stem cells. Stem Cell Reports 17, 1198–1214 (2022).

42. D. Pasini, A. P. Bracken, M. R. Jensen, E. Lazzerini Denchi, K. Helin, Suz12 is essential for mouse development and for EZH2 histone methyltransferase activity. EMBO J 23, 4061–4071 (2004).

43. R. Cao, Y. Zhang, SUZ12 is required for both the histone methyltransferase activity and the silencing function of the EED-EZH2 complex. Mol Cell 15, 57–67 (2004).

44. J. B. Suzich, A. R. Cliffe, Strength in diversity: Understanding the pathways to herpes simplex virus reactivation. Virology 522, 81–91 (2018).

45. A. Piunti, A. Shilatifard, The roles of Polycomb repressive complexes in mammalian development and cancer. Nat Rev Mol Cell Biol 22, 326–345 (2021).

46. K. E. Connelly, E. C. Dykhuizen, Compositional and functional diversity of canonical PRC1 complexes in mammals. Biochim Biophys Acta Gene Regul Mech 1860, 233–245 (2017).

47. Z. Gao, et al., PCGF homologs, CBX proteins, and RYBP define functionally distinct PRC1 family complexes. Mol Cell 45, 344–356 (2012).

48. A. Lagarou et al., dKDM2 couples histone H2A ubiquitylation to histone H3 demethylation during Polycomb group silencing. Genes Dev 22, 2799–2810 (2008).

49. H. F. Moussa et al., Canonical PRC1 controls sequence-independent propagation of Polycomb-mediated gene silencing. Nat Commun 10, 1931 (2019).

50. G. van Mierlo, G. J. C. Veenstra, M. Vermeulen, H. Marks, The Complexity of PRC2 Subcomplexes. Trends Cell Biol 29, 660–671 (2019).

51. S. Cooper et al., Jarid2 binds mono-ubiquitylated H2A lysine 119 to mediate crosstalk between Polycomb complexes PRC1 and PRC2. Nat Commun 7, 13661 (2016).

52. V. Kasinath et al., JARID2 and AEBP2 regulate PRC2 in the presence of H2AK119ub1 and other histone modifications. Science 371 (2021).

53. S. Sanulli et al., Jarid2 Methylation via the PRC2 Complex Regulates H3K27me3 Deposition during Cell Differentiation. Mol Cell 57, 769–783 (2015).

54. J. Son, S. S. Shen, R. Margueron, D. Reinberg, Nucleosome-binding activities within JARID2 and EZH1 regulate the function of PRC2 on chromatin. Genes Dev 27, 2663–2677 (2013).

55. D. Al-Raawi et al., A novel form of JARID2 is required for differentiation in lineage-committed cells. EMBO J 38 (2019).

56. E. Glancy et al., PRC2.1- and PRC2.2-specific accessory proteins drive recruitment of different forms of canonical PRC1. Mol Cell 83, 1393–1411 e1397 (2023).

57. Z. L. Watson et al., In Vivo Knockdown of the Herpes Simplex Virus 1 Latency-Associated Transcript Reduces Reactivation from Latency. J Virol 92 (2018).

58. D. A. Leib et al., A deletion mutant of the latency-associated transcript of herpes simplex virus type 1 reactivates from the latent state with reduced frequency. J Virol 63, 2893–2900 (1989).

59. A. M. Farcas et al., KDM2B links the Polycomb Repressive Complex 1 (PRC1) to recognition of CpG islands. Elife 1, e00205 (2012).

60. J. C. Brown, High G+C Content of Herpes Simplex Virus DNA: Proposed Role in Protection Against Retrotransposon Insertion. Open Biochem J 1, 33–42 (2007).

61. T. Gunther et al., A comparative epigenome analysis of gammaherpesviruses suggests cis-acting sequence features as critical mediators of rapid polycomb recruitment. PLoS Pathog 15, e1007838 (2019).

62. N. G. Naik et al., Epigenetic factor siRNA screen during primary KSHV infection identifies novel host restriction factors for the lytic cycle of KSHV. PLoS Pathog 16, e1008268 (2020).

63. N. Dietrich et al., REST-mediated recruitment of polycomb repressor complexes in mammalian cells. PLoS Genet 8, e1002494 (2012).

64. T. Du, G. Zhou, S. Khan, H. Gu, B. Roizman, Disruption of HDAC/CoREST/REST repressor by dnREST reduces genome silencing and increases virulence of herpes simplex virus. Proc Natl Acad Sci U S A 107, 15904–15909 (2010).

65. G. Zhou, T. Du, B. Roizman, HSV carrying WT REST establishes latency but reactivates only if the synthesis of REST is suppressed. Proc Natl Acad Sci U S A 110, E498–506 (2013).

66. A. R. Cliffe et al., Neuronal Stress Pathway Mediating a Histone Methyl/Phospho Switch Is Required for Herpes Simplex Virus Reactivation. Cell Host Microbe 18, 649–658 (2015).

67. Y. J. Machida, Y. Machida, A. A. Vashisht, J. A. Wohlschlegel, A. Dutta, The deubiquitinating enzyme BAP1 regulates cell growth via interaction with HCF-1. J Biol Chem 284, 34179–34188 (2009).

68. M. E. Sowa, E. J. Bennett, S. P. Gygi, J. W. Harper, Defining the human deubiquitinating enzyme interaction landscape. Cell 138, 389–403 (2009).

69. S. Misaghi et al., Association of C-terminal ubiquitin hydrolase BRCA1-associated protein 1 with cell cycle regulator host cell factor 1. Mol Cell Biol 29, 2181–2192 (2009).

70. H. Yu et al., The ubiquitin carboxyl hydrolase BAP1 forms a ternary complex with YY1 and HCF-1 and is a critical regulator of gene expression. Mol Cell Biol 30, 5071–5085 (2010).

71. A. Forrester et al., Construction and properties of a mutant of herpes simplex virus type 1 with glycoprotein H coding sequences deleted. J Virol 66, 341–348 (1992).

72. L. Benboudjema, M. Mulvey, Y. Gao, S. W. Pimplikar, I. Mohr, Association of the herpes simplex virus type 1 Us11 gene product with the cellular kinesin light-chain-related protein PAT1 results in the redistribution of both polypeptides. J Virol 77, 9192–9203 (2003).

73. S. A. Malin, B. M. Davis, D. C. Molliver, Production of dissociated sensory neuron cultures and considerations for their use in studying neuronal function and plasticity. Nat Protoc 2, 152–160 (2007).

74. R. E. Riccio, S. J. Park, R. Longnecker, S. J. Kopp, Characterization of Sex Differences in Ocular Herpes Simplex Virus 1 Infection and Herpes Stromal Keratitis Pathogenesis of Wild-Type and Herpesvirus Entry Mediator Knockout Mice. mSphere 4 (2019).

75. S. A. Stewart et al., Lentivirus-delivered stable gene silencing by RNAi in primary cells. RNA 9, 493–501 (2003).

76. S. McFarlane et al., The histone chaperone HIRA promotes the induction of host innate immune defences in response to HSV-1 infection. PLoS Pathog 15, e1007667 (2019).

77. T. Alandijany et al., Distinct temporal roles for the promyelocytic leukaemia (PML) protein in the sequential regulation of intracellular host immunity to HSV-1 infection. PLoS Pathog 14, e1006769 (2018).

78. L. G. Burman, V. P. Mauro, Analysis of rRNA processing and translation in mammalian cells using a synthetic 18S rRNA expression system. Nucleic Acids Res 40, 8085–8098 (2012).

