## Supplementary figures and images for "Histone H2A Ubiquitination Mediates the Establishment of Reactivation-Competent HSV-1 Latent Infection"

### Supplemental Figure 1

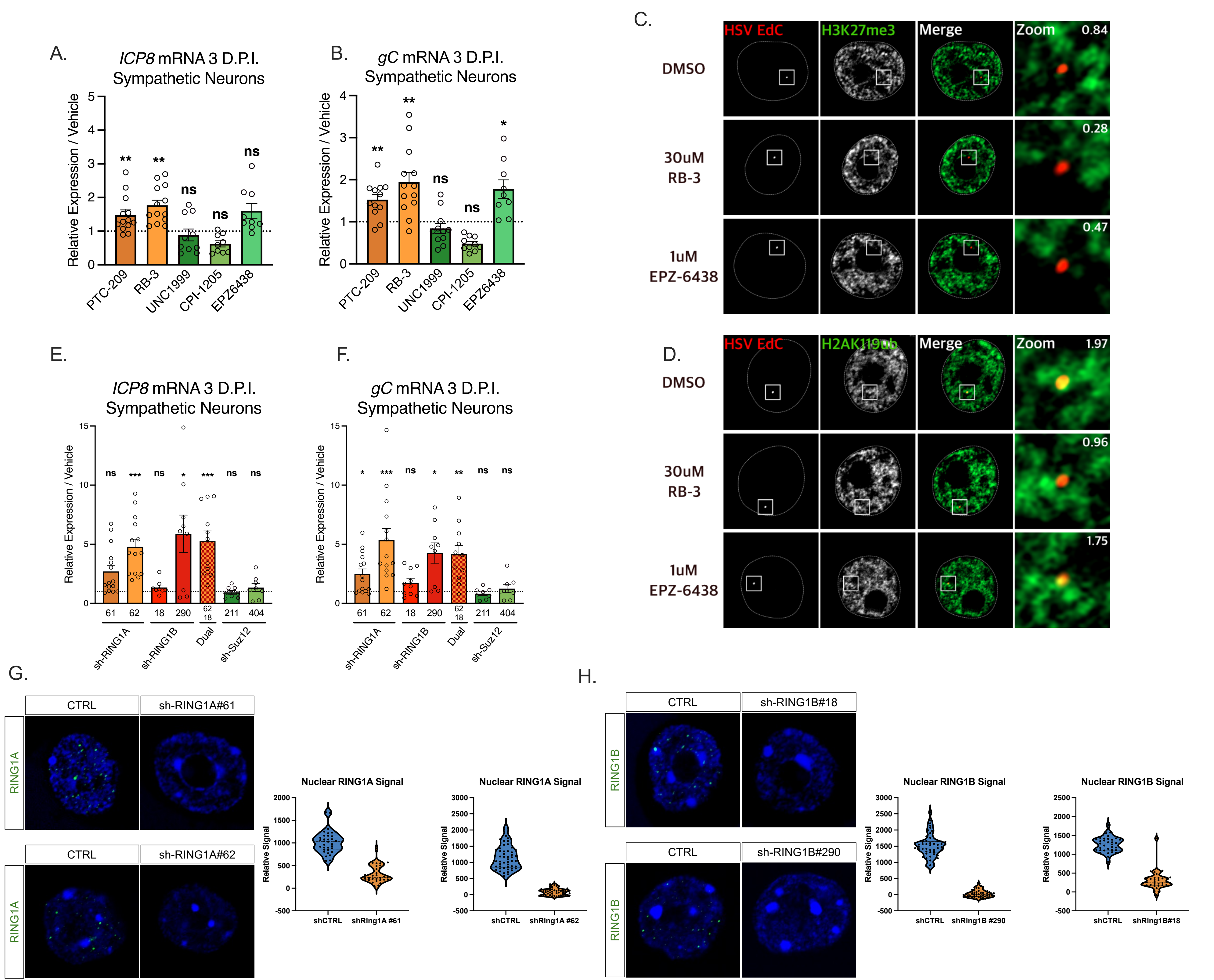
